# Enteroendocrine cells protect the stem cell niche by regulating crypt metabolism in response to nutrients

**DOI:** 10.1101/2022.10.01.508931

**Authors:** Heather A. McCauley, Anne Marie Riedman, Jacob R. Enriquez, Xinghao Zhang, Miki Watanabe-Chailland, J. Guillermo Sanchez, Daniel O. Kechele, Emily F. Paul, Kayle Riley, Courtney Burger, Richard A. Lang, James M. Wells

## Abstract

**Background and aims:** The intestinal stem cell niche is exquisitely sensitive to changes in diet, with high fat diet, caloric restriction, and fasting resulting in altered crypt metabolism and intestinal stem cell function. Unlike cells on the villus, cells in the crypt are not immediately exposed to the dynamically changing contents of the lumen. We hypothesized that enteroendocrine cells (EECs), which sense environmental cues and in response release hormones and metabolites, are essential for relaying the nutrient status of the animal to cells deep in the crypt.

**Methods:** We used the tamoxifen-inducible *VillinCreERT2* mouse model to deplete EECs (*Neurog3*^*fl/fl*^) from adult intestinal epithelium and we generated human intestinal organoids from wild-type and NEUROG3-null human pluripotent stem cells. We used indirect calorimetry, ^1^H-NMR metabolomics, mitochondrial live imaging, and the Seahorse bioanalyzer to assess metabolism. Intestinal stem cell activity was measured by proliferation and enteroid-forming capacity. Transcriptional changes were assessed using 10X Genomics single-cell sequencing.

**Results:** Loss of EECs resulted in increased energy expenditure in mice, an abundance of active mitochondria, and a shift of crypt metabolism to fatty acid oxidation. Crypts from mouse and human intestinal organoids lacking EECs displayed increased intestinal stem cell activity and failed to activate phospho-S6 ribosomal protein, a marker for activity of the master metabolic regulator mammalian target of rapamycin (mTOR). These phenotypes were similar to those observed when wild-type mice were deprived of nutrients.

**Conclusions:** Deletion of EECs recapitulated a fasting phenotype despite normal levels of ingested nutrients. These data suggest that EECs are required to relay nutritional information to the stem cell niche and are essential regulators of intestinal metabolism.

## Introduction

Oral nutrients are required to fuel all cellular and systemic functions throughout the body. The gastrointestinal tract digests and absorbs carbohydrates, proteins, fats, and micronutrients that then pass into the circulation to be used by other organs. However, as the initial site of nutrient sensing and nutrient absorption, the intestinal epithelium must first fuel itself to then be able to power the rest of the body. In fact, 20-25% of the body’s entire metabolic demand occurs within the intestine^1^.

The large metabolic demand of the mammalian small intestine is due in part to the fact that the epithelium is regenerated every ∼5 days from highly proliferative intestinal stem cells that reside in the crypts of Lieberkühn^2^. Intestinal stem cells give rise to daughter progenitor cells which go through several rounds of proliferation as they migrate upward through the transit amplifying zone while committing into the absorptive or secretory lineages in the terminally differentiated villus epithelium^2^. Some enteroendocrine cells and all Paneth cells instead migrate downward into the crypt region where a complex orchestra of secreted factors from epithelial and mesenchymal cells function as the intestinal stem cell niche^2^.

Enteroendocrine cells (EECs) of the gut represent the largest endocrine organ in the body, and one of their primary functions is to sense environmental cues like luminal nutrients and metabolites^3^. In response to these diverse inputs, EECs secrete over 20 distinct hormones, numerous other bioactive peptides, and small metabolites such as ATP^4^. The secreted products of EECs enter systemic circulation, interact with the enteric and central nervous system, and exert paracrine effects on local targets within the gut itself^3,4^. Despite their rarity, EECs are essential for several intestinal functions. Congenital loss of EECs results in failure to thrive and malabsorptive diarrhea upon ingestion of oral nutrition in humans^5^ and mouse^6,7^. We previously reported that EECs were required to maintain the electrophysiology supporting nutrient absorption in the small intestine^7^, placing EECs in a central role regulating basic cell biological functions of their neighboring enterocytes.

The diverse cellular functions that intestinal epithelial cells perform are supported by a range of cellular metabolic pathways^8^. Absorptive enterocytes are the most abundant cell type of the small intestine and primarily absorb nutrients and water. Macronutrient absorption and trafficking are energetically demanding processes, and as such enterocytes contain abundant mitochondria and largely utilize oxidative phosphorylation^8-10^. Like most proliferative zones in the body, intestinal crypts generate most of their energy via glycolysis^9-11^. However, healthy mitochondria, which perform numerous functions like reactive oxygen species production, cell death, and activation of biosynthetic pathways, in addition to energy production, are essential for proper intestinal stem cell function and differentiation into mature cell types^11-16^. Mitochondrial activity is higher in intestinal stem cells than in Paneth cells, and the metabolites lactate^13^ and pyruvate^11^ produced by glycolysis in Paneth cells have been demonstrated to serve as important niche factors for intestinal stem cell function. Moreover, loss of the mitochondrial chaperone protein *Hsp60*^12^ or of the transcription factor YY^16^, which is required for expression of mitochondrial complex I genes, resulted in loss of intestinal stem cell numbers and activity. Together, these studies demonstrate that while oxidative phosphorylation is not the dominant source of energy for the crypt as a whole, mitochondria are essential in maintaining intestinal stem cell function.

Crypt metabolism and intestinal stem cell function are exquisitely sensitive to changes in diet^17-21^. Intriguingly, when mice are fasted^21^ or fed a high-fat diet^17,18^, intestinal stem cells respond by upregulating mitochondrial fatty acid oxidation. In times of fat excess, the intestinal stem cell utilizes available nutrients to maximize efficiency of nutrient availability. Conversely, in times of nutrient deprivation, the intestinal stem cell shifts to fatty acid oxidation as a protective mechanism to preserve intestinal function^21^. Because cells deep within the crypt are far removed from the high nutritional environment of the lumen, we hypothesized that a nutrient-sensing cell such as the enteroendocrine cell may be involved in crypt adaptations to nutrient availability.

In this study, we identified a new role for EECs in regulating the cellular metabolism of the crypt. We found that acute loss of EECs increased mitochondrial activity and fatty acid oxidation in whole intestinal crypts, resulting in increased intestinal stem cell activity in mouse small intestine and human intestinal organoids. Loss of EECs recapitulated many aspects of fasting, including loss of activity of mammalian target of rapamycin (mTOR) signaling. mTOR signaling is the main downstream sensor of availability of nutrients, guiding stem cells in many organs to adjust to nutritional availability. Together, our data suggest that EECs are required to maintain intestinal stem cell homeostasis.

## Results

### Conditional loss of EECs alters whole-body and intestinal metabolism within days

Congenital loss of EECs results in malabsorptive diarrhea with poor postnatal survival^6,7^, so to circumvent this obstacle we crossed *Neurog3*^*fl/fl*^ animals with the tamoxifen-inducible *VillinCreERT2*^22^ to deplete EECs in adult mice (Supplemental Figure 1). We used the tdTomato Ai9 reporter allele^23^ to monitor for Cre recombination efficiency (Figure 1A). 10 days after tamoxifen administration, EECs were diminished from the mid-jejunum by 96%, with a corresponding reduction in hormone transcripts and proteins (Figure 1A-B and Supplemental Figure 1A-D). At 10 days, EEC-deficient mice displayed no outward signs of poor health, no significant change in weight, and fecal pellets were well-formed. However, upon internal examination, the jejunum, ileum, cecum, and proximal colon were distended and filled with liquid contents in EEC-deficient mice (Supplemental Figure 1E). We reasoned that the poor efficiency of the *Villin* promotor in the distal colon^24^ (Supplemental Figure 1F) allowed for adequate water resorption and the formation of solid feces despite the proximal liquid enteral contents. The gross phenotype of the small bowel suggested that loss of EECs disrupted intestinal physiology and function in the adult animal.

**Figure 1:**
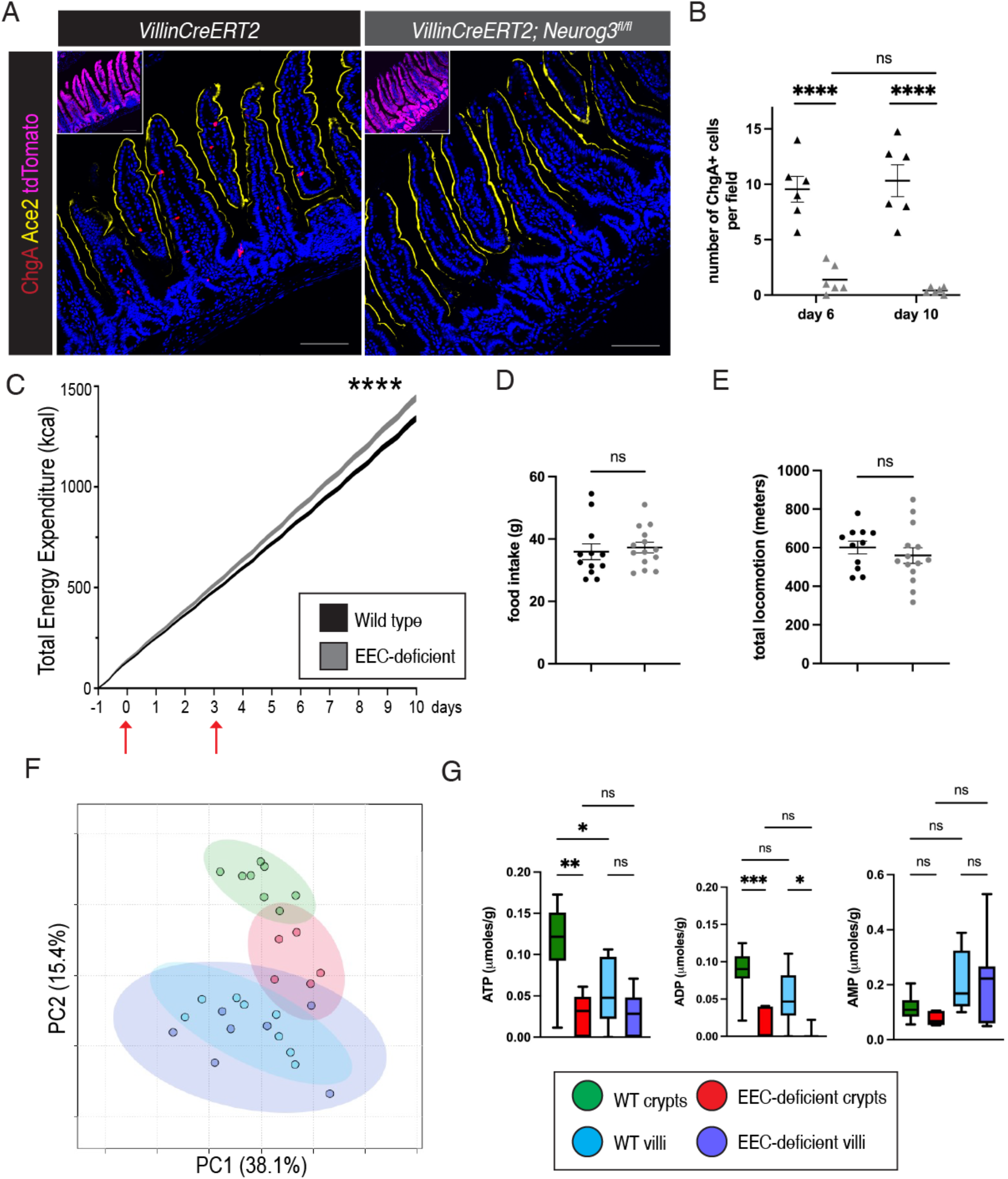
EECs are required to maintain whole-body and intestinal metabolism. A. Immunofluorescence staining for Chromogranin A (ChgA) in *VillinCreERT2;Neurog3*^*fl/fl*^*;tdTomato* and *VillinCreERT2;tdTomato* wild-type control jejunum 6 days after tamoxifen administration. Scale bars = 100μm. B. Quantification of the representative image shown in F (day 6, **** *P*<.0001; day 10, **** *P*<.0001; no difference between day 6 and day 10). n=6 mice per genotype per time point. Statistical significance determined using 2-way ANOVA with Tukey’s multiple comparison’s test. C. Total energy expenditure over time measured by whole-body respirometry. Red arrows indicate tamoxifen injections. n=12 wild-type, n=14 EEC-deficient mice. **** *P*<.0001; statistical significance determined using simple linear regression. D. Total food eaten by wild-type and EEC-deficient animals as measured by metabolic cages. n=12 wild-type, n=14 EEC-deficient mice. E. Total activity performed by wild-type and EEC-deficient animals as measured by metabolic cages. n=12 wild-type, n=14 EEC-deficient mice. F. Principal component analysis (PCA) scores plot, PC1 (38.1% EV) and PC2 (15.4% EV), indicating the presence of unique metabolic profiles in crypts and villi from wild-type and EEC-deficient animals. Green: wild-type crypt (n=8); red: EEC-deficient crypt (n=5); light blue: WT villi (n=8); purple: EEC-deficient villi (n=7). G. Normalized concentrations of ATP and ADP per gram of tissue. Significance calculated by one-way ANOVA with Tukey’s multiple comparison’s test. ATP, \**P =* 0.0231, ** *P* = 0.0029, ** *P* = 0.0019; ADP, * *P* =0.0113, *** *P* =0.0006, **** *P* <.00001.

EECs have a well-established role in regulating many aspects of systemic metabolism. To determine whether loss of EECs impacted whole-body metabolism, wild-type and *VillinCreERT2;Neurog3*^*fl/fl*^ animals were housed in metabolic cages for assessment by indirect calorimetry (Figure 1C). During the period of acclimation, there was no difference in any parameter measured between genotypes; however, approximately one day following the first dose of tamoxifen, *VillinCreERT2;Neurog3*^*fl/fl*^ animals began to increase their total energy expenditure over wild-type animals (Figure 1C). This difference in energy expenditure, calculated by the abbreviated Weir equation as an additive function of oxygen consumption and carbon dioxide production, continued to increase for the course of the 10-day experiment. There was no difference in food consumed or activity between wild-type and EEC-deficient groups (Figure 1D-E), suggesting the increased energy expenditure occurred primarily at the cellular level.

Considering that the intestine is a major metabolic organ, consuming approximately 20-25% of the body’s total oxygen requirements^1^, we questioned whether loss of EECs would impact cellular metabolism of intestinal epithelial cells directly. We performed ^1^H-NMR metabolomic profiling on epithelial cells isolated from wild-type and EEC-deficient jejunum (Figure 1F). Because there are well-defined metabolic differences in cellular metabolism between crypts and villi^9^, we separated these compartments for our analysis. Thirty-four unique metabolites were identified and quantified (Supplemental Figure 2) and a principal component analysis (PCA) showed significant overlap between wild-type and EEC-deficient villus samples. However, EEC-deficient crypts clustered distinctly from wild-type crypts, clustering closer to the villus samples (Figure 1F). Two of the significant metabolites altered in EEC-deficient crypts compared to wild-type were ATP and ADP (Figure 1G). In wild-type animals, ATP was higher in the crypt than in the villus, consistent with the glycolytic nature of the highly proliferative crypt compartment^9^. Loss of EECs resulted in significant reduction of both ATP and ADP in the crypt compared to wild-type crypts (Figure 1G). In EEC-deficient samples, crypt ATP was reduced to villus levels, whereas ADP was very low or not detected at all (Figure 1G). These data suggest that EECs are required for normal cellular metabolism along the crypt-villus axis, and that in the absence of EECs, the crypt adopts a metabolic profile more closely resembling that of the villus.

### EECs regulate mitochondrial activity in intestinal crypts

EEC-deficient crypts adopted a more villus-like metabolic profile (Figure 1F), which predominantly utilize mitochondrial oxidative phosphorylation to meet energetic demands^9^. To investigate this further we assessed mitochondrial activity using a live-imaging approach using tetramethylrhodamine methyl ester (TMRM), a fluorescent readout of mitochondrial activity (Figure 2A). Cells in wild-type villi had high mitochondrial activity, and loss of EECs had no obvious impact on villus mitochondrial activity, consistent with the ^1^H-NMR profiling (Figure 1F). In comparison to the villus preparations, wild-type crypts had low mitochondrial activity as determined by few TMRM+ cells. However, EEC-deficient crypts displayed abundant TMRM+ cells, indicating highly active mitochondria in nearly every crypt cell (Figure 2A). TMRM+ active mitochondria were observed in mice which had undergone tamoxifen-induced depletion of EECs 28 days prior to analysis (Supplemental Figure 3A), suggesting that the metabolic reprogramming of the intestine was a stable phenotype. Activation of TMRM+ mitochondria did not occur immediately upon loss of *Neurog3*, as 24 hours post-tamoxifen administration there was no difference between genotypes, but robust TMRM+ activity was observed 3 days after tamoxifen administration (Supplemental Figure 3A). This increase in mitochondrial activity was not due an increase in mitochondrial content since there was no change in gene expression of mitochondrial inner and outer membrane proteins nor protein expression of outer mitochondrial membrane protein Tom20 between genotypes (Supplemental Figure 3B-C). These data suggested that EECs regulate mitochondrial activity in the crypt.

**Figure 2:**
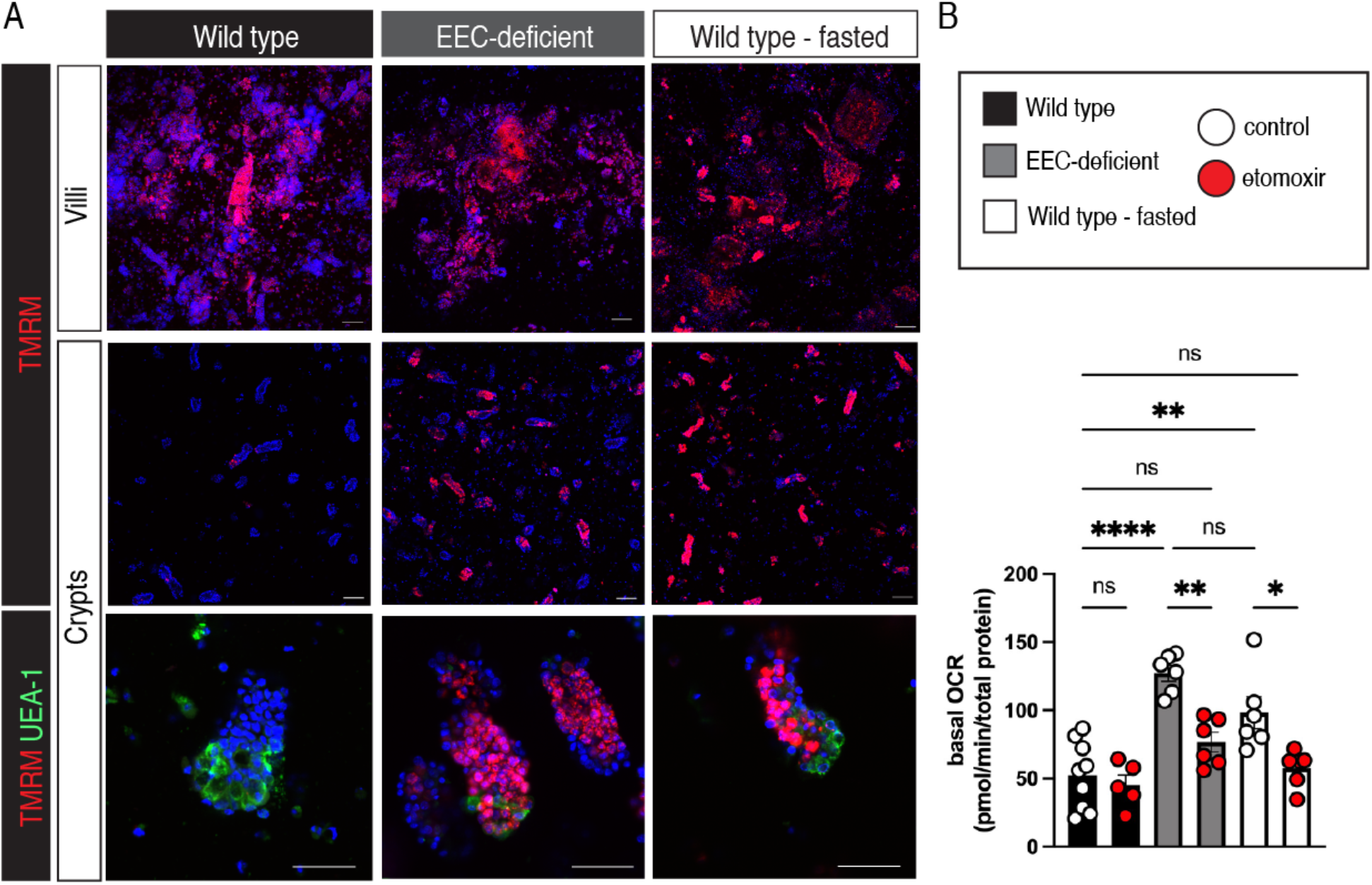
EECs regulate mitochondrial activity in intestinal crypts. A. Crypts and villi were separated from jejunum of wild-type animals, EEC-deficient animals, and wild-type animals which were fasted overnight and incubated in TMRM and Hoescht to visualize active mitochondria via live confocal microscopy. Bottom panels display zoomed-in representative crypts with Paneth cells labeled with UEA-1. Scale bars = 10μm (top), 50μm (bottom). B. Crypts of WT and EEC-deficient animals were plated for Seahorse analysis of mitochondrial activity. Basal oxygen consumption of wild-type and EEC-deficient crypts, with and without the fatty acid oxidation inhibitor etomoxir (**** *P* .00001, WT v KO; * *P* = .018, WT v WT Fasting; ** *P =* .005, KO v KO + etomoxir). Statistics calculated using one-way ANOVA with Tukey’s multiple comparison’s test. Error bars represent SEM.

Cells of the crypt can dynamically alter their metabolism in response to changes in nutrient availability. For example, when mice are fasted, the crypt responds by activating mitochondrial fatty acid metabolism to maintain homeostasis in the absence of nutrients^21^. We observed abundant TMRM+ active mitochondria in the crypts of wild-type mice which had been fasted overnight, mimicking the elevated mitochondrial activity observed in EEC-deficient crypts (Figure 2A). To functionally investigate whether increased TMRM fluorescence correlated with fatty acid oxidation and oxidative phosphorylation, we measured crypt mitochondrial activity using the Seahorse bioanalyzer (Figure 2B). We observed significantly increased basal oxygen consumption in EEC-deficient crypts and in fasted wild-type crypts, consistent with increased mitochondrial activity, compared to fed wild-type crypts. We next cultured crypts in etomoxir, a chemical inhibitor that blocks the entry of fatty acids into mitochondria for oxidation. Wild type crypts had low levels of basal oxygen consumption, and this was unaffected by treatment with etomoxir. In contrast, the high basal oxygen consumption in EEC-deficient crypts and fasted wild-type crypts were restored to wild-type fed levels by etomoxir (Figure 2B). This indicated that the elevated mitochondrial activity observed in the absence of nutrients or in the absence of nutrient-sensing EECs was due to activation of mitochondrial fatty acid oxidation.

### EECs regulate intestinal stem and progenitor activity

Crypt metabolism is a major determinant of intestinal stem cell function^9,25^. In particular, fasting increases proliferation and enhances intestinal stem cell activity^21^. We therefore investigated whether the altered crypt metabolism in EEC-deficient animals impacted intestinal stem and progenitor activity. We induced deletion of EECs in 8–12-week-old animals and monitored cell proliferation and ISC marker expression 10 days after the first tamoxifen injection (Figure 3A-B). EEC-deficient crypts had elevated cell proliferation and expanded expression of the stem and progenitor marker Olfm4^26^, consistent with fasted wild-type animals (Figure 3A-B). There was with no change in apoptosis between genotypes (Supplemental Figure 4). To determine if the increases in proliferation and ISC markers in the EEC-deficient animals translated into increased ISC activity, we quantified enteroid-forming capacity of isolated crypts grown in culture. EEC-deficient crypts had 2-3 fold higher enteroid-forming capacity than wild type, consistent with fasted crypts^21^ (Figure 3C).

**Figure 3:**
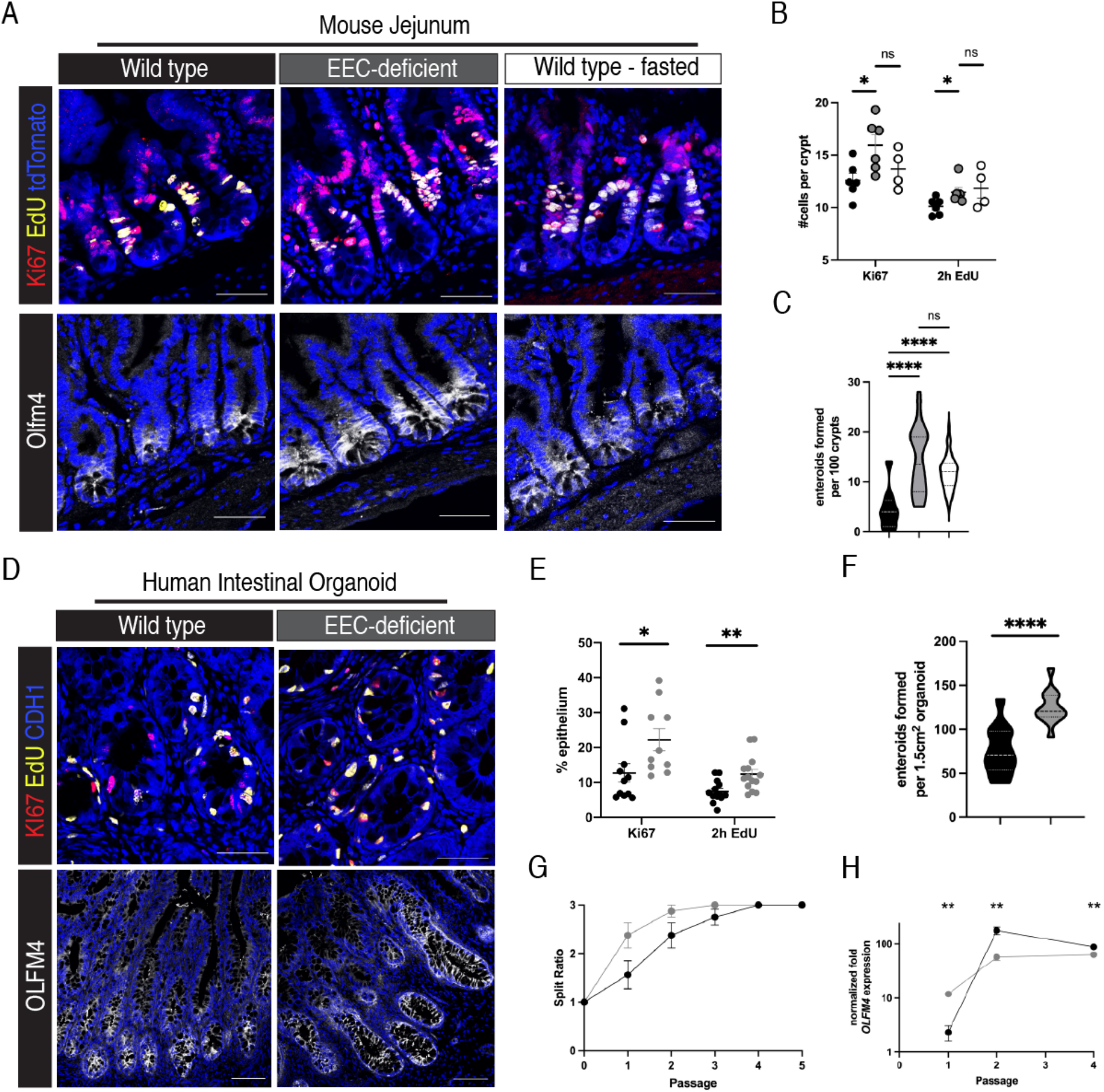
EECs regulate intestinal stem and progenitor activity. A. Immunofluorescence staining for Ki67, EdU (2h pulse), and Olfm4 in wild-type, EEC-deficient, and fasted wild-type mouse jejunal crypts. DAPI counterstains nuclei in blue. Scale bars = 50μm. B. Quantification of proliferation in (A). * *P =* 0.02, Ki67; * *P =* 0.04, EdU. Statistics calculated using unpaired t-test with the two-stage step-up (Benjamini, Krieger, and Yekutiele method). C. Enteroid forming capacity of wild-type, EEC-deficient, and fasted wild-type mouse jejunal crypts (**** *P*<.0001). n=18 wells from 3 wild type mice, 24 wells from 4 EEC-deficient mice, and 48 wells from 4 fasted wild-type mice. Statistics calculated using ordinary one-way ANOVA with Tukey’s multiple comparison’s test. D. Immunofluorescence staining for Ki67, EdU (2h pulse), and OLFM4 in wild-type and EEC-deficient human intestinal organoids. DAPI counterstains nuclei in blue. Scale bars = 50μm (top panels), 20μm (bottom panels). E. Quantification of proliferation in (C). * *P*=.032, Ki67, ** *P*=.007, EdU. Statistics calculated using unpaired t-test with the Holm-Sidak method. F. Enteroid forming capacity of wild-type and EEC-deficient human intestinal organoid crypts (**** *P*<.0001). n=12 wells from two independent organoids per genotype. Statistics calculated using unpaired, two-tailed t-test G. Visualization of HIO-derived enteroid growth as depicted by split ratios at each passage. The slope of the curves was significantly (* *P*=0.03) different from passage 0 to passage 2. Statistical significance determined by simple linear regression. H. qPCR depicting normalized mRNA expression for *OLFM4* in HIO-derived enteroids collected at passages 1, 2, and 4. Statistics calculated by unpaired, two-tailed t-test.

The functional role of EECs in regulating human intestinal stem cell activity is unknown. We therefore performed similar analyses of human intestinal organoids lacking EECs. We generated EEC-deficient pluripotent stem cell-derived human intestinal organoids as previously described^7^. Organoids were transplanted into mice for an additional 12-weeks of growth. This results in intestinal tissues with robust crypt-villus architecture containing all mature cell types, including intestinal stem cells^27^. Like the murine model, EEC-deficient organoids displayed increased proliferation and expanded OLFM4+ stem and progenitor cells (Figure 3D-E). Moreover, crypts isolated from organoid transplants had increased enteroid-forming capacity compared to wild-type (Figure 3F).

Crypts from EEC-deficient intestine had a phenotype resembling that of a nutrient-deficient environment despite the fact that animals were ingesting normal levels of food. From this, we hypothesized that EECs couple nutrient availability with crypt behaviors. Therefore, we predicted that if we eliminated the need for EECs to transmit the nutrient status to the crypt, ISC behavior would not be impacted by the absence of EECs. To do this, we grew crypts for multiple passages in culture where intestinal stem cells were consistently exposed to high levels of nutrients. While EEC-deficient enteroids had a growth advantage immediately upon culture generation, requiring to be split earlier than wild-type cultures, this difference normalized within two passages (Figure 3G). Expression of OLFM4 also normalized between genotypes over multiple passages. At passage 1, OLFM4 levels were ∼12-fold higher in EEC-deficient enteroids (Supplemental Figure 4) but by passage 4 expression had increased to ∼60-fold over passage 1 levels and had largely normalized between genotypes alongside the normalization in growth rate (Figure 3G-H). These data suggested that EECs are essential for relaying the nutrient status to the crypt *in vivo* when the crypt is restricted from direct access to nutritional cues, and that intestinal stem cells require input from the EEC to maintain homeostasis *in vivo*.

### Without EECs, intestinal stem and progenitor cells upregulate lipid metabolism genes and downregulate master nutrient sensor mTORC1

To further investigate the mechanism by which EECs regulate metabolism in the intestinal crypt, we performed single-cell RNA (scRNA) sequencing of transplanted human intestinal organoids with and without EECs. After filtering and quality control of three wild-type organoids and one EEC-deficient organoid, 14,968 cells were recovered from the pooled wild-type samples and 4853 cells were recovered from the EEC-deficient sample. As previously described^28^, scRNA-sequencing of wild-type organoids identified all the expected epithelial cell types, including intestinal stem cells, proliferative progenitors, Paneth-like cells, enterocytes, goblet cells, and EECs (Supplemental Figure 5A-B), whereas organoids generated from NEUROG3-null pluripotent stem cells were specifically deficient in EECs (Supplemental Figure 5A). We performed gene ontology (GO) analysis on the differentially expressed genes between wild-type and EEC-deficient organoids in multiple cell clusters. Strikingly, in the stem and progenitor cell clusters, we observed that all 20 of the top biological processes upregulated in EEC-deficient organoids were associated with cellular metabolism, especially of lipids (Figure 4A-B). Lipid metabolism was also upregulated in Paneth cells in EEC-deficient organoids (Figure 4B). Examples of genes involved in lipid metabolism that were significantly upregulated in EEC-deficient crypt cells include arylacetamide deacetylase (*AADAC)*, alpha ketoreductase family members 1C1 and 1C3 (*AKR1C1, AKR1C3*), apolipoproteins A1 and A4 (*APOA1, APOA4*), catalase (*CAT*), fatty acid binding protein 2 *(FABP2)*, and microsomal triglyceride transfer protein *(MTTP)* (Figure 4C). These human data are consistent with the functional data obtained in mouse, and together all data suggest that loss of EECs results in upregulation of fatty acid oxidation and a deficit of nutrient sensing in the crypts.

**Figure 4:**
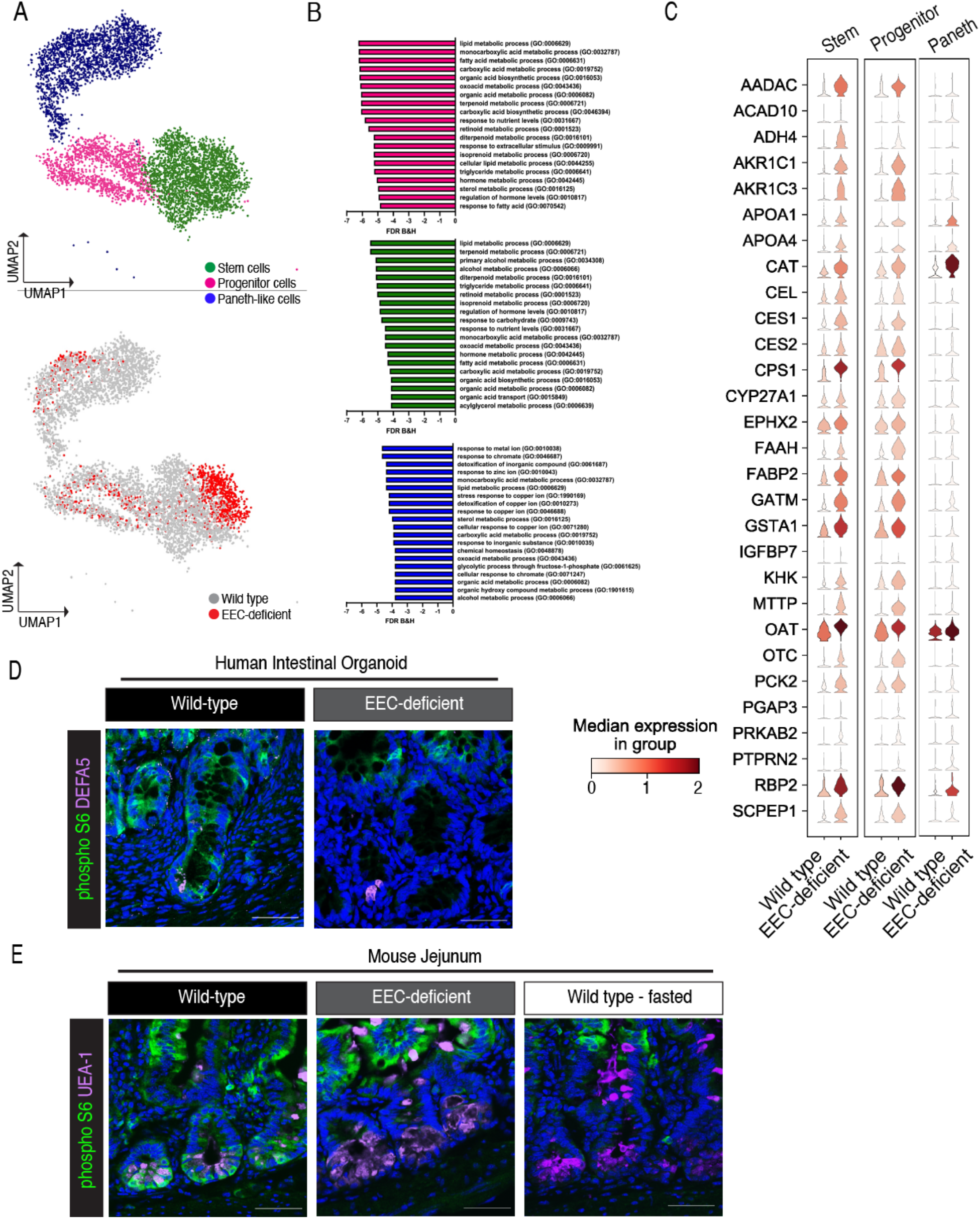
Without EECs, intestinal stem and progenitor cells upregulate lipid metabolism genes and downregulate master nutrient sensor mTORC1. A. Transplanted human intestinal organoids were dissociated into single-cells and sequenced using the 10X Genomics platform. UMAP depicting the stem, progenitor, and Paneth-like cell clusters of the integrated dataset colored by cluster (top) and colored by genotype (bottom). B. Gene ontology analysis of the differentially expressed genes in EEC-deficient stem (top, green), progenitor (middle, pink), and Paneth-like (bottom, blue) cell clusters. The top 20 biological processes are displayed. C. Violin plots for a selection of genes represented in GO:0006629, lipid metabolic process, in the stem, progenitor, and Paneth-like cell clusters in wild-type and EEC-deficient human intestinal organoids. D. Immunofluorescence staining for pS6 in crypts of wild-type and EEC-deficient human intestinal organoids. DEFA5 marks Paneth cells. DAPI counterstains nuclei in blue. Scale bar = 50μm. E. Immunofluorescence staining for pS6 in small intestinal crypts of wild-type, EEC-deficient, and fasted wild-type mice. UEA-1 marks Paneth cells. DAPI counterstains nuclei in blue. Scale bar = 50μm.

As a final investigation of how crypts from EEC-deficient small intestine might be impaired in their ability to sense nutrients, we analyzed activity of the master cellular pathway governing nutrient sensing, mammalian target of rapamycin (mTOR). In response to abundant nutrients, mTOR complex 1 (mTORC1) signaling, as measured by the phosphorylation of downstream target S6 kinase, activates an array of downstream catabolic pathways. In fasted animals, when nutrients are low, mTORC1 signaling is decreased in the crypt^19^. Similar to the fasted state, crypts in EEC-deficient mice and human intestinal organoids show a marked reduction of phosphorylated S6 in the crypts compared to fed wild-type animals (Figure 4D-E). These data suggest that EECs are required to transmit nutritional information to activate the nutrient-sensing, master metabolic regulator mTORC1 in the crypt.

## Discussion

In this study, we uncovered a novel role for enteroendocrine cells in the regulation of crypt metabolism and intestinal stem cell homeostasis. By using mice and human pluripotent stem cell-derived intestinal organoids, we found that EECs relay the nutrient status of the gut to the crypt, essential for controlling intestinal stem cell activity and activating the master metabolic regulator mTORC1. In the absence of EECs, intestinal crypts adopt fasting-like characteristics, including increased mitochondrial activity, activation of fatty acid oxidation, increased proliferation and increased intestinal stem cell activity. These functional analyses were supported by single-cell sequencing results in human intestinal organoids which demonstrated that metabolic pathways in intestinal stem and progenitor cells were profoundly impacted by loss of EECs. Moreover, loss of EECs diminished the activity of the master nutrient-sensing mTORC1 pathway, phenocopying wild-type mice which had been restricted from food. All data suggest that that despite the presence of nutrients, crypts from EEC-deficient animals behaved as if they were in a state of nutrient deprivation.

We found that loss of EECs results in an altered metabolic environment in all cells residing in the crypt, including intestinal stem cells, proliferative progenitor cells, and Paneth cells. Paneth cells are important niche cells contributing metabolites and growth factors to the intestinal stem cell to maintain stemness and function^2^ and maintain an independent metabolic identity from intestinal stem cells^13^. Moreover, mTORC1 signaling is known to be high in Paneth cells and acts as a key regulatory pathway coupling nutrient availability to intestinal stem cell function^19^. Under normal conditions, Paneth cells mainly use glycolysis; however, in EEC-deficient human intestinal organoids, we observed increased expression of lipid metabolism genes. This shift from glycolysis to lipid metabolism correlated with reduced mTORC1 signaling in human and mouse Paneth cells. These disruptions in Paneth cell metabolism raise the possibility that EECs may lie upstream of Paneth cells in regulating niche homeostasis.

EEC regulation of intestinal metabolism has implications for numerous gastrointestinal pathologies, including colorectal cancer, gastric cancer, and inflammatory bowel disease (IBD). Most cancers alter their cellular metabolism to consume high levels of glucose and produce ATP rapidly via aerobic glycolysis, termed the “Warburg effect.” Additionally, a number of other metabolic pathways, including glutamine, lipid, oxidative phosphorylation, and one-carbon metabolism, can all be dysregulated in cancer, including colorectal cancer^29^. Mitochondrial dysfunction is a hallmark of IBD, partially attributed to genetic susceptibility factors and largely due to the microbial-epithelial environment of the gut epithelium itself^8^. In Crohn’s disease and ulcerative colitis, many studies have demonstrated diminished mitochondrial energy production, increase in reactive oxygen species (ROS), increase in cell stress, and diminished autophagy – all driving an inflammatory disease process^8^. The pathological changes in the intestine following loss of EECs are relevant to several disease contexts. For example, one of the most highly upregulated genes we identified in Paneth cells from human intestinal organoids which lacked EECs was *catalase*, an antioxidant enzyme which protects cells from oxidative stress. In addition, gene ontology analysis suggested that Paneth cells from EEC-deficient organoids may have altered metal ion processing, including that of zinc, copper, and chromate. These metals can be oxidized to produce free radicals and contribute to ROS-induced stress throughout the body, and are pathogenic contributors to gastrointestinal cancers and IBD^30^. Our data suggest that EECs may serve a yet unidentified role in protecting Paneth cells from oxidative stress.

EECs secrete over 20 distinct biologically active peptides and other small molecules, such as ATP, in response to environmental stimuli^3^, and EEC number and hormone production are altered in diseases such as IBD and type 2 diabetes. EECs are overall increased in mouse models of colitis and human patients with ulcerative colitis and Crohn’s disease, although the populations of hormones are skewed in each disease^31^. In ulcerative colitis, there is a significant increase in serotonin (5-HT), but no change in peptide YY, pancreatic polypeptide, GLP-1, or somatostatin-expressing cells. Conversely, in patients with Crohn’s disease, there is a similar significant increase in 5-HT-expressing cells, increased GLP-1, no change in somatostatin, and loss of peptide YY and pancreatic polypeptide compared to healthy controls. While there have been limited reports of EEC activity in human patients with colorectal cancer, there have been some *in vitro* studies reported. In colon cancer cell lines, ghrelin^32,33^ stimulated proliferation and metastatic potential, whereas activation of the GLP-1 receptor inhibited cell growth and induced apoptosis in tumor-bearing mice^34^. Moreover, the shift in cellular metabolism in response to changing nutrient availability within the tumor environment^29^ suggests potential roles for multiple nutrient-sensing EECs in tumor proliferation and pathogenesis.

In moving forward, it will be important to determine how EECs integrate particular nutritional or microbial cues with specific secreted products to integrate luminal inputs with metabolic pathways and functional outcomes. With this information, it should be possible to develop practical new therapeutic avenues for patients with metabolic disease, IBD, or gastrointestinal cancers.

## Materials and Methods

### Mice

B6.Cg-Tg(Vil1-cre/ERT2)23Syr/J ^22^ (JAX# 020282), B6.Cg-*Gt(ROSA)26Sor*^*tm9(CAG-tdTomato)Hze*^*/J* ^23^ (JAX# 007909; Ai9), and *Neurog3*^*fl/fl* 6^ were maintained on a C56Bl/6 background. At 8-12 weeks of age, animals were administered two doses of tamoxifen (25mg/kg), dissolved in corn oil, 3 days apart. For all experiments, controls considered “wild type” were *VillinCreERT2;Neurog3*^*+/+*^ animals administered tamoxifen following the same protocol as *VillinCreERT2;Neurog3*^*fl/fl*^ mice. Mice with and without the tdTomato reporter were used interchangeably. Animals were housed in a specific pathogen free barrier facility in accordance with NIH Guidelines for the Care and Use of Laboratory Animals. All experiments were approved by the Cincinnati Children’s Hospital Research Foundation Institutional Animal Care and Use Committee (IACUC2022-0002) and carried out using standard procedures. Mice were maintained on a 12-hour light/dark cycle and had ad libitum access to standard chow and water. Mice were housed at 22ºC at 30-70% humidity.

### Diarrhea score

The abdomen was opened and visually inspected for the consistency of intestinal contents. Animals were scored based on luminal distension and the liquidity of luminal contents in the small intestine, cecum, and proximal colon. Animals were given a score of 0 for no liquid contents and well-defined stool pellets forming along the colon; a score of 1 for no liquid contents and poorly defined stool pellets merging together in the colon; a score of 2 for mild distension, some liquid contents in the small intestine and/or cecum, and poorly defined stool pellets merging together in the colon; a score of 3 for severe distension, watery luminal contents in the small intestine and/or cecum, and poorly defined stool pellets merged together in the proximal colon.

### Serum hormone levels

Hormone concentrations in the mouse serum were determined by using Milliplex™ Mouse Metabolic Hormone Expanded Panel – Metabolism Multiplex Assay (MilliporeSigma, #MMHE-44K) according to manufacturer’s protocol.

### Indirect Calorimetry

8 to 12-week-old male mice were moved from standard housing to metabolic cages (Promethion, Sable Systems International) for 14 days according to standard protocol. Mice were acclimatized for 3 days prior to administration of the first dose of tamoxifen. Mice were continuously recorded at ambient room temperature (22°C) on a 14L:10D light cycle with the following measurements taken every 5 minutes using Sable Systems International acquisition software (Promethion Live v.21.0.3): oxygen consumption (VO_2_), carbon dioxide emission (VCO_2_), energy expenditure (EE), respiratory exchange ratio (RER), food intake, water intake, and spontaneous locomotor activity (cm s^-1^) in the XY plane. Food and water were available *ad libitum*. Energy expenditure was calculated using the abbreviated Weir formula. Respiratory exchange ratio (RER) was calculated by the ratio VCO_2_/VO_2_. Mass dependent variables (VO_2_, VCO_2_, energy expenditure) were not normalized to body weight. Food and water intake were measured by top-fixed load cell sensors, from which food and water containers were suspended into the sealed cage environment. Raw data were exported using Sable Systems International ExpeData software v.1.9.27 and analyzed using Sable Systems International MacroInterpreter Software v.2.43. Reported data represent a combination of two independent experiments conducted separately.

### Crypt/villus separation

The small intestine was flushed with ice-cold PBS without Mg^2+^ or Cl^2+^ (Gibco) or with DMEM (Gibco) containing 10% FBS (Sigma-Aldrich) and divided into duodenum, jejunum, and ileum. The jejunum was filleted open and rinsed vigorously to remove mucus and debris, then cut into 1-2cm segments and transferred to a 15mL conical containing 5mL 200mM EDTA (Invitrogen) in PBS without Mg^2+^ or Cl^2+^. Tissue segments were shaken vigorously by hand for 30 seconds then rotated for 30 minutes at 4C. Tissue segments were then transferred to a new 15mL conical containing 5mL 1.5% sucrose and 1% sorbitol in PBS without Mg^2+^ or Cl^2+^ and shaken by hand for 2 minutes at a rate of 2 shakes per second to dislodge crypts. The solution containing crypts and villi was then filtered through a 70um filter set atop a 50mL conical tube, with crypts enriched in the filtered fraction. Villi were collected by washing the top of the filter with additional PBS without Mg^2+^ or Cl^2+^ and recovering the unfiltered fraction into a new conical tube. All steps were performed on ice. Crypts and villi were inspected via microscopy then centrifuged at 150g for 10 minutes at 4C prior to additional assays.

### Immunofluorescence

Mouse intestine and human intestinal organoids were fixed in 4% paraformaldehyde, cryopreserved in 30% sucrose, embedded in OCT, and frozen at -80C until cryosectioned. 8μm cryosections were mounted on Superfrost Plus slides and permeabilized, blocked and stained according to standard protocol. Primary antibodies used are listed in the table below. All secondary antibodies were conjugated to Alexa Fluor 488, 546/555/568 or 647 (Invitrogen) and used at 1:500 dilution. EdU was retrieved and stained using the Click-It EdU Alexa Flour 647 Imaging Kit (Invitrogen). Images were acquired using a Nikon A1 GaAsP LUNV inverted confocal microscope and NIS Elements software (Nikon). For images used for quantification, at least three 20X images were taken with well-oriented crypt-villus architecture and the average counts from the three images represented. On average, 30-40 crypts per animal were counted.

| Antibody | Company | Catalog # | Host | Dilution |
| --- | --- | --- | --- | --- |
| Ace2 | R&D Systems | AF3437 | Goat | 1:200 |
| Chromogranin A | ImmunoStar | 20085 | Rabbit | 1:200 |
| Carbonic Anhydrase 4 | R&D Systems | AF2414 | Goat | 1:200 |
| Ki67 | Cell Marque | 275R-14 | Rabbit | 1:200 |
| Cleaved Caspase-3 | Cell Signaling Technology | 9661S | Rabbit | 1:200 |
| DEFA5 | Novus Biologicals | NB110-60002 | Mouse | 1:60,000 |
| Olfm4 (mouse specific) | Cell Signaling Technology | 39141S | Rabbit | 1:200 |
| OLFM4 | Cell Signaling Technology | 14369T | Rabbit | 1:200 |
| Phospho-S6 ribosomal protein (S235/236) | Cell Signaling Technology | 2211S | Rabbit | 1:200 |
| UEA-1, DyLight 649 | Vector Laboratories | DL-1068 | n/a | 1:1,000 |
| UEA-1, fluorescein | Invitrogen | L32476 | n/a | 1:1,000 |

### qPCR

RNA was extracted using Nucleospin RNA extraction kit (Macharey-Nagel) and reverse transcribed into cDNA using Superscript VILO (Invitrogen) according to manufacturer’s instruction. qPCR was performed using Quantitect SYBR® Green PCR kit (QIAGEN). For mitochondrial genes, a custom Taqman array pre-loaded with the mouse mitochondrial panel (Applied Biosystems) was used with Taqman Fast Advanced Master mix. SYBR Green and Taqman qPCR were run on a QuantStudio 3 Flex Real-Time PCR System (Applied Biosystems). Relative expression was determined using the ΔΔCt method and normalizing to 18S (mouse) or CPHA (human). Primers used are listed in the table below.

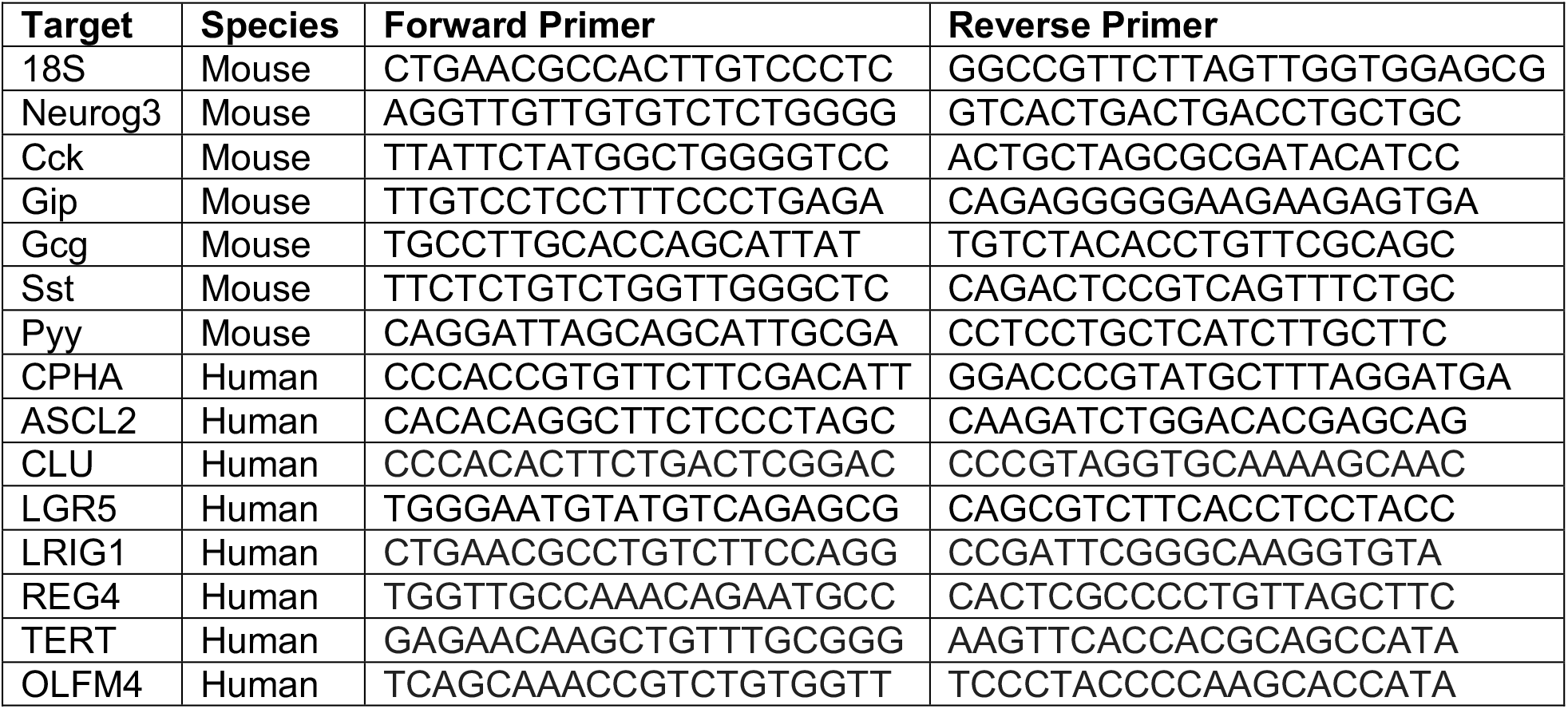

### Human Intestinal Organoids

Wild-type (H1) and EEC-deficient (NEUROG3^-/-^) human pluripotent stem cells^35,36^ were maintained, differentiated into intestinal organoids, and transplanted into the kidney capsule of immunocompromised NOD.Cg-*Prkdc*^*scid*^ *Il2rg*^*tm1Wjl*^/SzJ mice (JAX #005557) as previously described^7,37^. Organoids were harvested after 10-14 weeks of growth. Organoids were opened and washed to remove mucus, dead cells, and cellular debris from the lumen, then equally sized ∼1.5cm^2^ segments containing well-developed crypt-villus architecture were placed in a 15mL conical tube containing 5mL TryPLE Dissociation Reagent (GIBCO) for 30 minutes, shaking at 4C. Organoids were vigorously shaken to dislodge villi then moved to a sylgard-coated dish where crypts were scraped from the serosal tissue using the back of curved forceps. Crypts and villi were inspected via microscopy then centrifuged at 150g for 10 minutes at 4C prior to additional assays.

### TMRM

Crypt and villus fractions were suspended in Live Cell Imaging Solution (Invitrogen) containing 100nM tetramethylrhodamine methyl ester (TMRM) (Invitrogen), 10ug/mL Hoescht (Thermo Scientific), and Ulex Europaeus Agglutinin I (UEA I) conjugated to DyLight 649 (Vector Laboratories) or fluorescein (Invitrogen), both at 1:1000 dilution. Samples were incubated for 30 minutes at 37C, washed with PBS, centrifuged at 150g for 5 minutes, resuspended in phenol-red-free, growth-factor-reduced basement-membrane Matrigel (Corning #356231), and plated on glass-bottom dishes for confocal live-imaging. Once the Matrigel had solidified, Live Cell Imaging Solution was overlaid on the samples. Images were acquired using a Nikon A1 GaAsP LUNV inverted confocal microscope and NIS Elements software (Nikon). Laser power was set based on TMRM intensity in wild-type villus samples and not adjusted between samples.

### Seahorse

Mouse jejunal crypts were analyzed using the Seahorse XF96e bioanalyzer (Agilent Technologies) according to manufacturer’s instructions and according to protocols adapted from others^38,39^. The day before the assay, the 96 well plate was coated with basement membrane Matrigel (Corning) diluted 1:10 in complete Seahorse medium and set overnight at room temperature. The day of the assay, the coated plate was incubated in a non-CO_2_ incubator at 37C for 2 hours prior to plating crypts. 500 crypts per well were plated in complete Seahorse medium and allowed to attach for 1 hour at 37C in a non-CO_2_ incubator prior to analysis. Some wells were injected with etomoxir (4μM final concentration). After the experiment was completed, total protein was calculated using a Pierce BCA protein assay (Thermo Scientific) and all values normalized to total protein.

### Enteroid formation assay

Mouse jejunal crypts were plated at a density of 100 crypts per well in a 24-well plate in phenol-red-free, growth-factor-reduced basement-membrane Matrigel (Corning #356231) and fed with enteroid growth media containing 50% L-WRN^40^. 72 hours after plating, the number of spherical enteroids growing in each well were counted.

Human intestinal organoids were dissociated as described above. The villus fraction was discarded. Crypts were split equally into 6 wells in a 24-well plate in phenol-red-free, growth-factor-reduced basement-membrane Matrigel (Corning #356231) and fed with human IntestiCult Organoid Growth Media (STEMCELL technologies). 96 hours after plating, the number of spherical enteroids growing in each well were counted.

### NMR

Crypt and villus fractions were separated as above then snap-frozen and stored at -80C until sample processing. The modified Bligh and Dyer extraction^41^ was used to obtain polar metabolites from the crypt and villus. The samples were homogenized for 30 s at 5000 rpm in cold methanol and water with 2.8mm ceramic beads. The cold chloroform and water were added to the samples to bring the final methanol:chloroform:water ratio to be 2:2:1.8. The polar phase was dried by vacuum and resuspended in 600 uL of NMR buffer containing 100mM phosphate buffer, pH7.3, 1mM TMSP (3-Trimethylsilyl 2,2,3,3-d4 propinoate), 1mg/mL sodium azide, prepared in D_2_O. NMR experiments were acquired on a Bruker Avance III HD 600 MHz spectrometer with 5mm, BBO Prodigy probe (Bruker Analytik, Rheinstetten, Germany). All data were collected at a calibrated temperature of 298 K using the noesygppr1d pulse sequence and processed using Topspin 3.6 software (Bruker Analytik). For a representative sample, two-dimensional data 2D ^1^H-^13^C heteronuclear single quantum coherence (HSQC) were collected to assist with metabolites assignment. Total of 34 polar metabolites were assigned and quantified using Chenomx® NMR Suite profiling software (Chenomx Inc. version 8.4) based on the internal standard, TMSP. The concentrations were then normalized to the tissue weights prior to all statistical analyses and log transformation and mean centering data scaling were used for multivariate analysis. All the metabolomics data analysis were performed using R studio and MetaboAnalyst^42^.

### Western blot

Protein was extracted from crypt fractions in ice-cold RIPA buffer (Thermo Scientific) with 0.5mM EDTA and Halt Protease and Phosphatase inhibitor cocktail (100X, Thermo Scientific), sonicated, and the supernatant quantified by Pierce BCA assay (Thermo Scientific) before 1:1 dilution in Laemmle buffer containing 2-mercaptoethanol. Protein gels were run according to standard procedure on a 4-12% Bolt Bis-Tris Plus gel (Invitrogen), blocked with Intercept Blocking Buffer (LI-COR), and imaged using an Odyssey LI-COR CLx imaging system.

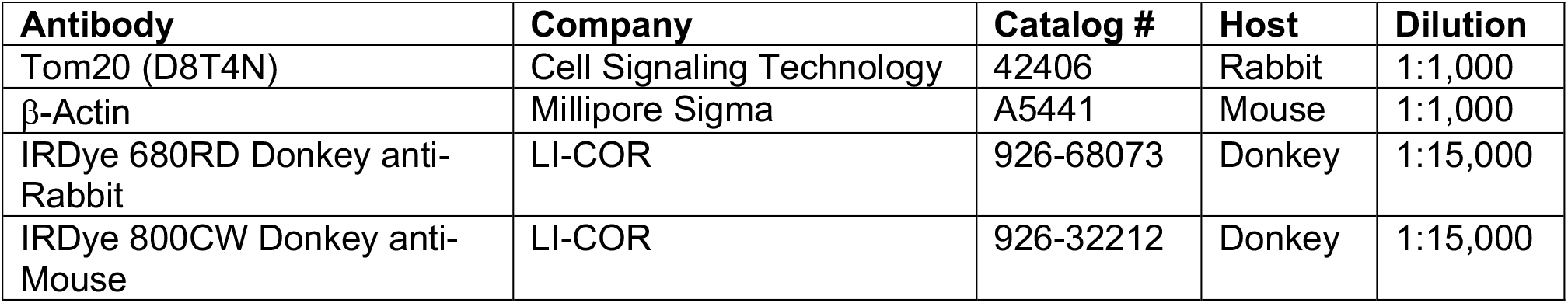

### Single-cell RNA-Sequencing

Human intestinal organoids were dissociated and the villus and crypt fractions from each organoid were pooled together. After centrifugation, cells were incubated in 0.05% Trypsin-EDTA (Thermo Fisher Scientific) for 10 minutes at 37C while shaking, washed with 10% FBS, vigorously pipetted to create a single-cell suspension, then filtered through a 100μm and 40μm filter. Filtered cells were centrifuged at 500g for 5 minutes at 4C, resuspended in 5% FBS + 0.5% BSA + 10mm Y-27632 (Tocris) in PBS and sorted for viability by exclusion of Sytox Blue dead cell stain (Thermo Fisher Scientific) on a BD FACS Aria II. Live cells were collected in 5% FBS + 0.5% BSA in PBS, adjusted to a concentration of 10^6^ cells/mL. Single cell RNA-Seq library preparation was performed by the CCHMC Single Cell Genomics Core using the Chromium 3’v3 GEX Kit (10x Genomics). Approximately 12,800 cells were loaded to achieve 8,000 captured cells per sample to be sequenced. Sequencing was performed by the CCHMC DNA Sequencing core using the NovaSeq 6000 (Illumina) sequencing platform with an S2 flow cell to obtain approximately 320 million reads per sample. Raw scRNA-seq data was converted to FASTQ files and then aligned to the human genome [hg19] using CellRanger v3.0.2 (10x Genomics).

scRNA-seq data were analyzed using python 3.7.3 via Scanpy (v1.7.0). After default basic filtering, quality control of samples included cells with maximum 8000 genes and with less than 40 percent mitochondrial related genes. Default parameters were used for leiden clustering, visualization via UMAP, and cluster identification via differential gene expression. Differentially expressed gene lists were generated for each cluster with thresholds of logfold change greater than 1 and a Wilcoxon adjusted p value less than 0.05. Gene lists were loaded into ToppGene Suite^43^ to assess overlap with annotated gene ontology lists. Parameters used were FDR B&H with a cutoff of 0.05 and *P* value was calculated using hypergeometric probability mass function.

## Acknowledgements

We would like to acknowledge the assistance of the ^1^H-NMR Metabolomics Core in the Division of Pathology and Laboratory Medicine, the Research Flow Cytometry Core in the Division of Rheumatology, and the Single Cell Genomics Core in the Division of Developmental Biology at Cincinnati Children’s Hospital Medical Center.

---

**Supplemental Figure 1:**
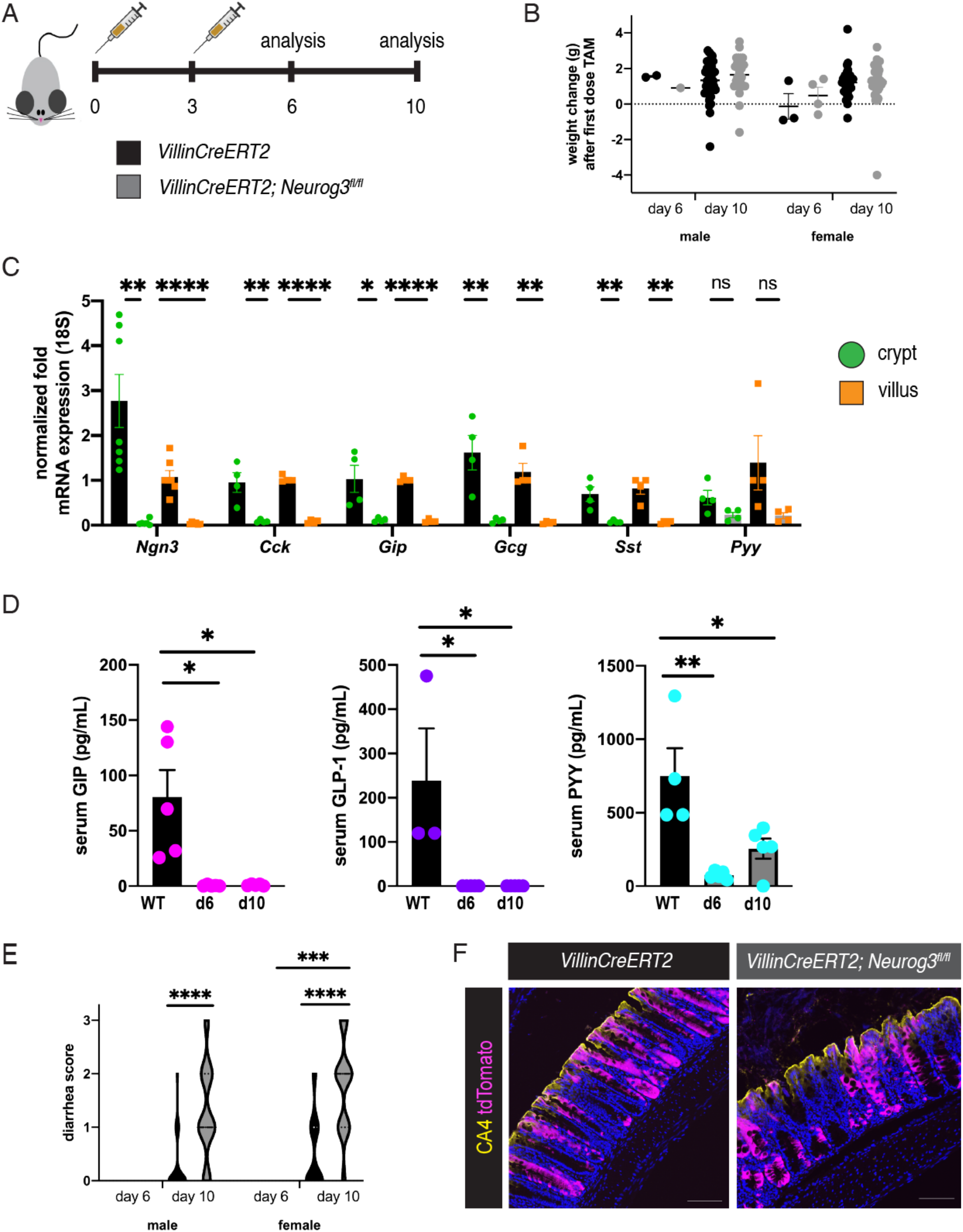
Enteroendocrine cells are lost in adult *VillinCreERT2* intestine within 10 days. A. Schematic of tamoxifen (TAM) dosing strategy. Mice aged 8 weeks or older were administered two doses of tamoxifen (25mg/kg) 3 days apart. Animals were harvested 6 days or 10 days after the initial dose to evaluate loss of enteroendocrine cells (EECs). Age-matched, non-littermate *VillinCreERT2* mice (black bars) were used as controls. Some animals also carried the Ai9 *Rosa26-*^*flox-STOP-flox-tdTomato*^ reporter allele to evaluate the efficiency of recombination. B. Body weight of animals dosed with tamoxifen. Statistical significance determined by 2-way ANOVA. C. Diarrhea score of *VillinCreERT2;Neurog3*^*fl/fl*^ animals (gray bars) compared to *VillinCreERT2* wild-type animals (black bars) dosed with tamoxifen (male, \*\*\*\**P*<.0001; female, **** *P*<.0001, \*\*\**P*=.0003 change from day 6 to day 10). Notably, many *VillinCreERT2* wild type animals exhibited mild diarrhea 10 days after tamoxifen treatment, although this did not reach significance over animals analyzed at 6 days which never displayed diarrheal symptoms. Statistical significance determined by 2-way ANOVA. D. tdTomato expression in the colon of wild-type *VillinCreERT2;tdTomato* and *VillinCreERT2;Neurog3*^*fl/fl*^*;tdTomato* animals 10 days after tamoxifen administration. Carbonic anhydrase 4 (CA4) marks the surface epithelial cells of colonocytes. Scale bars = 100μm. E. Jejunum of *VillinCreERT2* and *VillinCreERT2;Neurog3*^*fl/fl*^ animals were harvested 6 days after initial tamoxifen treatment, separated into crypt (green circles) and villus (orange squares) compartments, and analyzed for expression of enteroendocrine-specific genes by qPCR. *Ngn3* (crypt, \*\**P*=.001405; villus, \*\*\*\**P*=.000029), *Cck* (crypt, \*\**P*=.007638; villus, \*\*\*\**P*<.000001), *Gip* (crypt, \**P*=.02405; villus, \*\*\*\**P*<.000001), *Gcg* (crypt, \*\**P*=.007729; villus, \*\**P*=.001133), and *Sst* (crypt, \*\**P*=.008389; villus, \*\**P*=.001345), *Pyy* (crypt, *P*=.071144; villus, *P*=.099997). Statistical significance determined using the Holm-Sidak method. F. Serum levels of GIP, GLP-1, and PYY 6-10 days after tamoxifen administration (GIP, day 6, * *P*=.0108, day 10, * *P*=.0114; GLP-1, day 6, * *P*=.0329, day 10, * *P*=.0329; PYY, day 6, ** *P*=.0012, day 10, * *P*=.0141). Statistical significance determined using 1-way ANOVA with Tukey’s multiple comparison’s test.

**Supplemental Figure 2:**
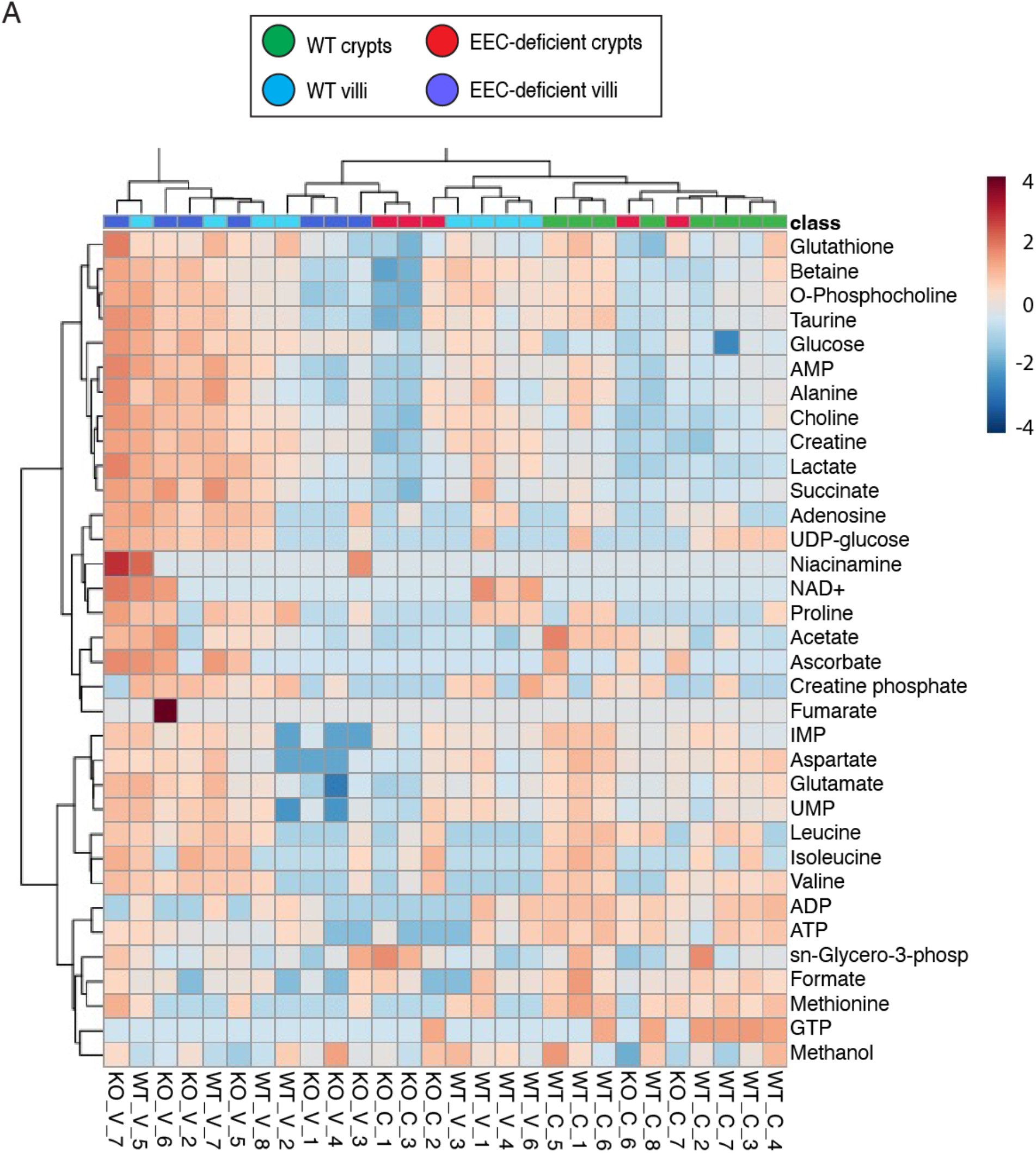
^1^NMR-Metabolomic profiling of WT and EEC-deficient small intestine. A. Ward eucidean hierarchical clustering heatmap depicting the 34 metabolites identified by ^1^H-NMR Metabolomics across the wild-type and EEC-deficient crypt (C) and villus (V) samples.

**Supplemental Figure 3:**
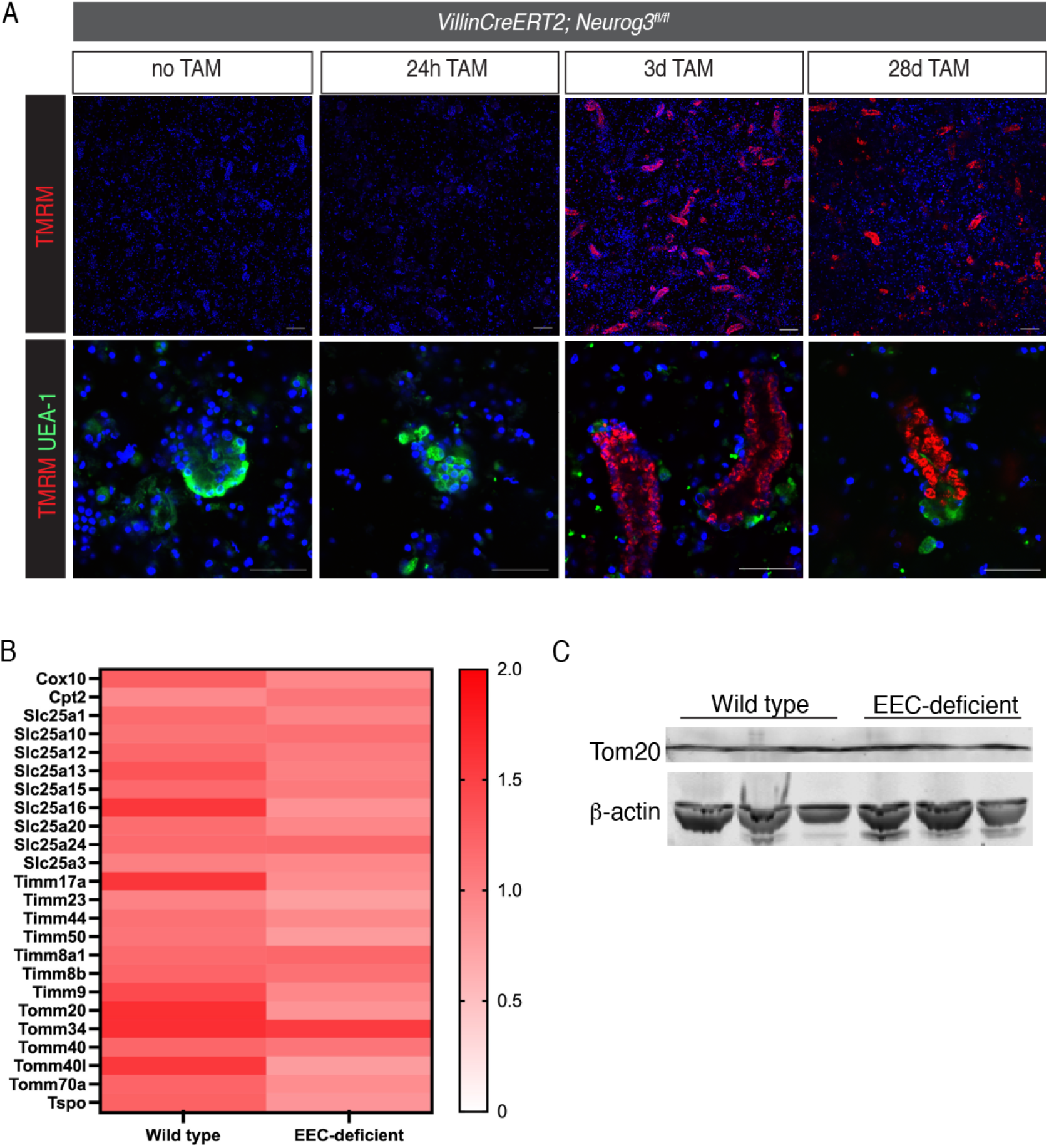
Tamoxifen induces mitochondrial activity by 3 days and does not change total mitochondrial content. A. Live confocal microscopy of TMRM-labeled mitochondria in crypts isolated from *VillinCreERT2; Neurog3fl/fl* animals not treated with tamoxifen (TAM), 24 hours, 3 days, or 28 days after tamoxifen administration. Bottom panels display zoomed-in representative crypts with Paneth cells labeled with UEA-1. Hoescht labels nuclei in blue. Scale bars = 10μm (top), 50μm (bottom). B. Taqman array for genes encoding inner and outer mitochondrial membrane proteins in wild-type and EEC-deficient crypts. Average expression of 3 animals per genotype is depicted in the heatmap, normalized to 18S. C. Crypt cells were probed for mitochondrial membrane protein Tom20 by Western blot. β-actin was used as a loading control. n=3 animals per genotype.

**Supplemental Figure 4:**
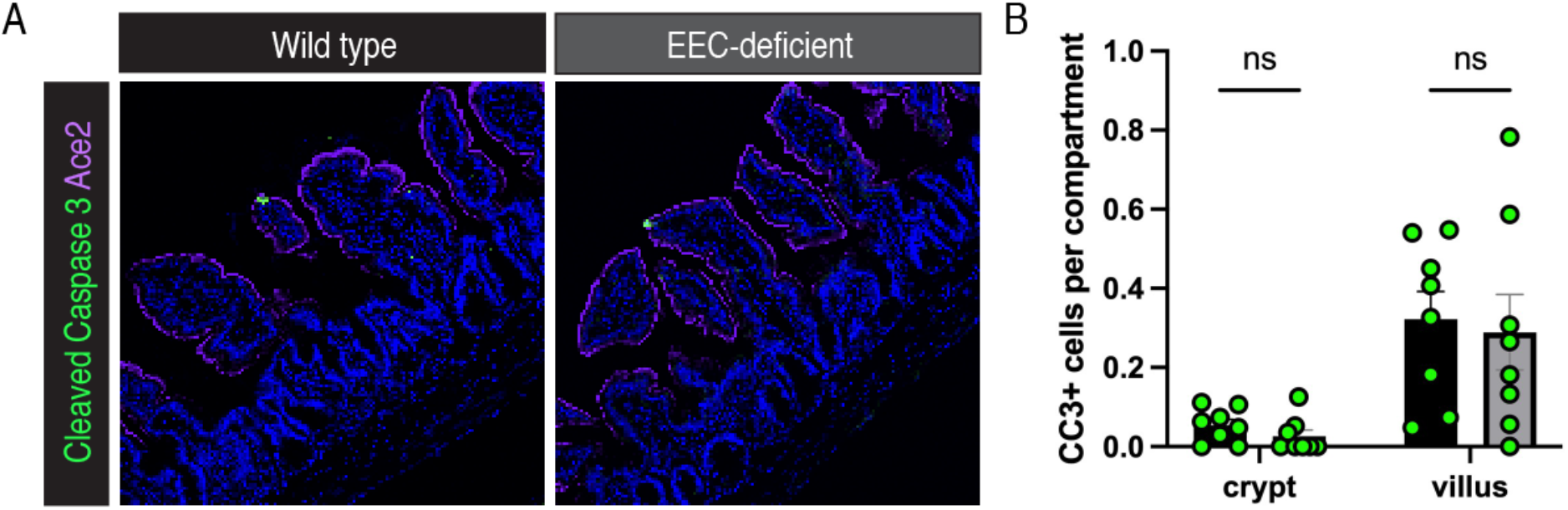
Loss of EECs does not impact apoptosis in mouse small intestine. A. Immunofluorescence staining for cleaved caspase-3 in wild-type and EEC-deficient small intestine. Ace2 marks the brush border and DAPI counterstains nuclei. Scale bars = 50μm. B. Quantification of (A). Three well-oriented images per mouse were averaged and the number of cleaved caspase-3-positive cells per number of crypts and villi per image represented. n=8 animals per genotype.

**Supplemental Figure 5:**
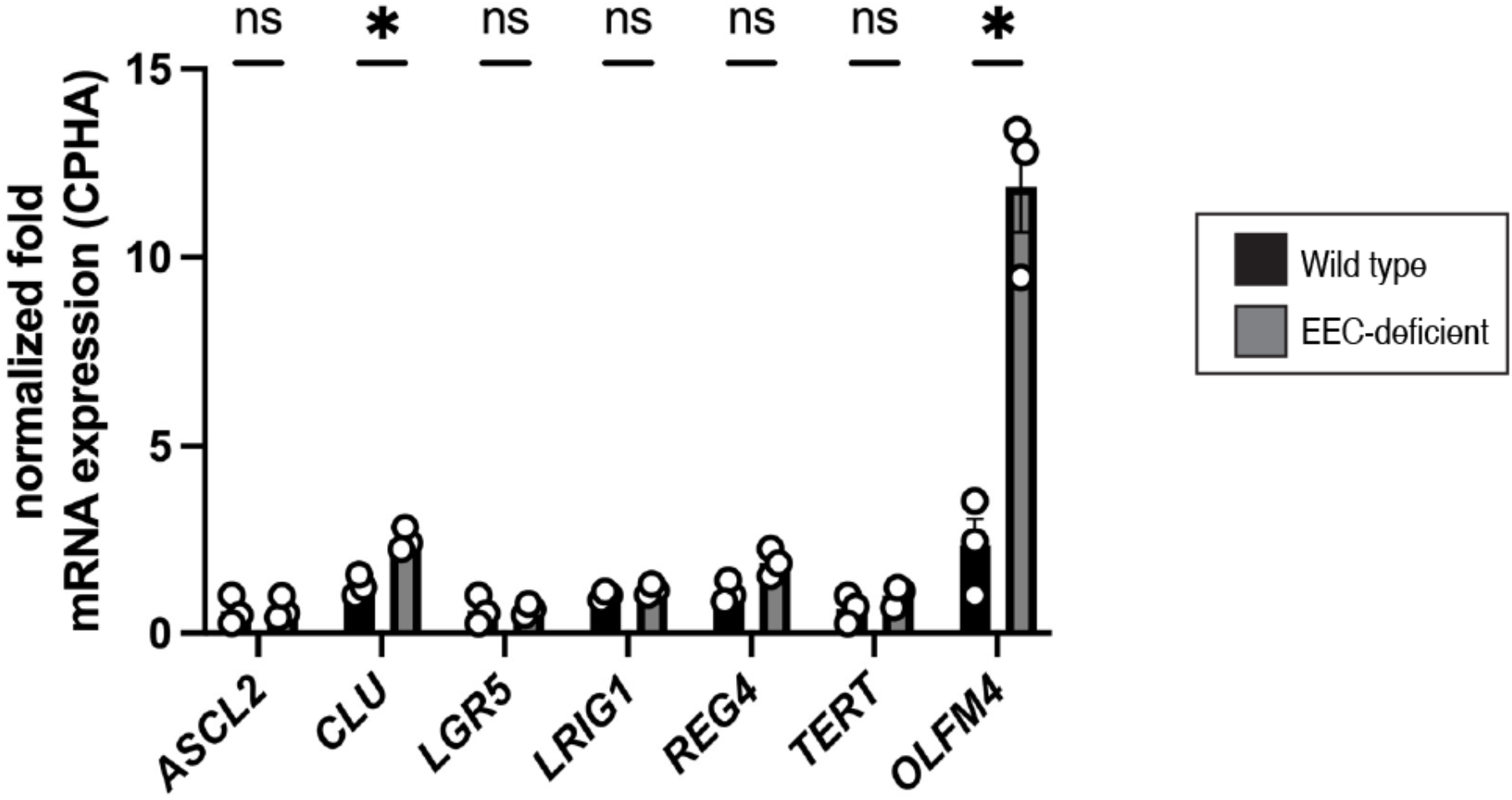
EEC-deficient enteroids derived from human intestinal organoids upregulate *OLFM4* at first passage. A. Normalized mRNA expression for a selection of intestinal stem cell genes at first passage of enteroids derived from wild-type and EEC-deficient human intestinal organoids. Statistics calculated by unpaired, two-tailed t-test.

**Supplemental Figure 6:**
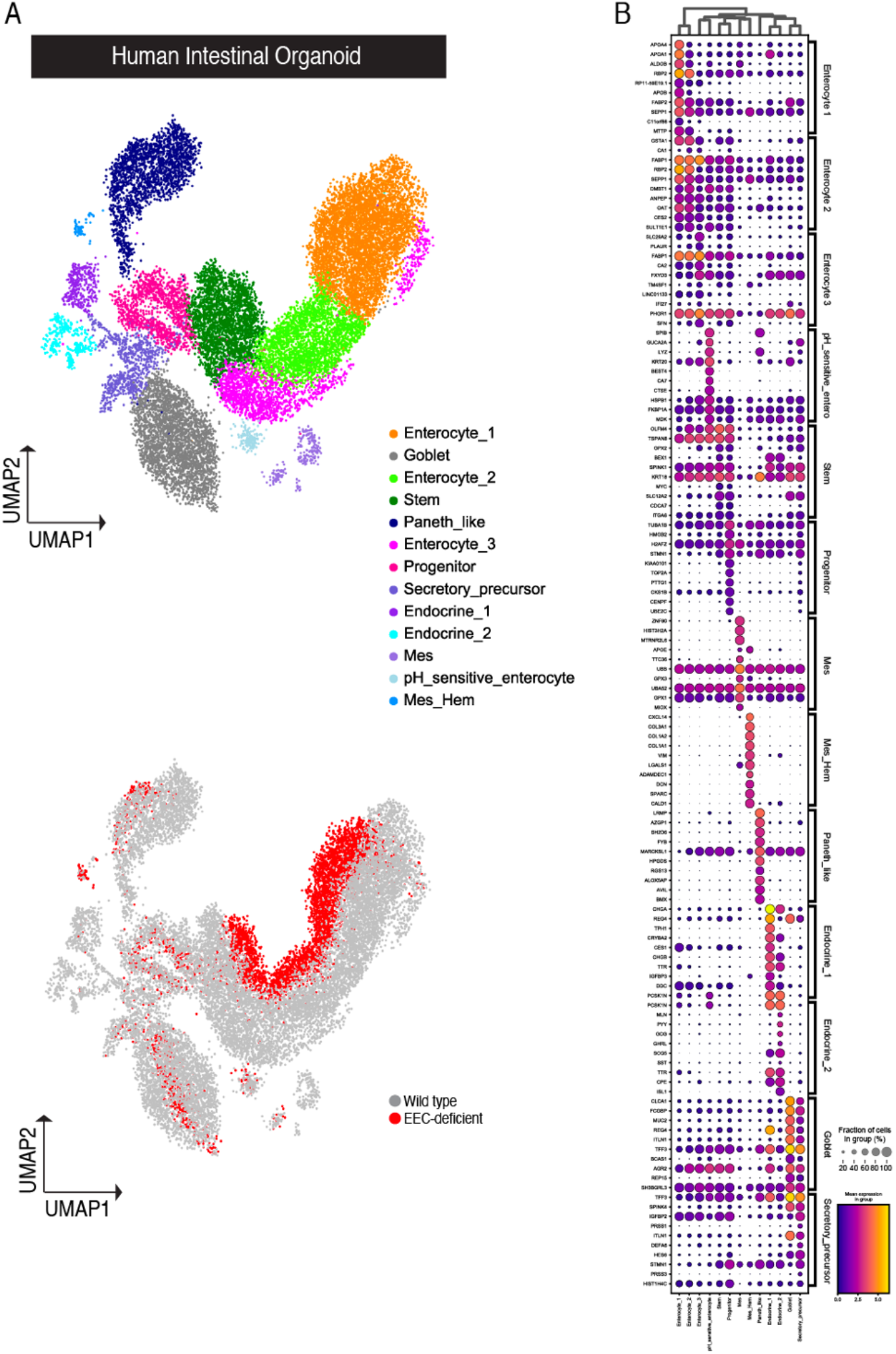
Human intestinal organoids generate expected cell populations by single-cell sequencing. A. Transplanted human intestinal organoids were dissociated into single-cells and sequenced using the 10X Genomics platform (n=3 wild-type, n=1 EEC-deficient). UMAP depicting the integrated dataset of all cells colored by cluster (top) and colored by genotype (bottom). B. Dot plot depicting the expression of the top 10 genes in each cell cluster in the integrated dataset.

